# Critical Assessment of Metagenome Interpretation – a benchmark of computational metagenomics software

**DOI:** 10.1101/099127

**Authors:** Alexander Sczyrba, Peter Hofmann, Peter Belmann, David Koslicki, Stefan Janssen, Johannes Dröge, Ivan Gregor, Stephan Majda, Jessika Fiedler, Eik Dahms, Andreas Bremges, Adrian Fritz, Ruben Garrido-Oter, Tue Sparholt Jørgensen, Nicole Shapiro, Philip D. Blood, Alexey Gurevich, Yang Bai, Dmitrij Turaev, Matthew Z. DeMaere, Rayan Chikhi, Niranjan Nagarajan, Christopher Quince, Fernando Meyer, Monika Balvoit, Lars Hestbjerg Hansen, Søren J. Sørensen, Burton K. H. Chia, Bertrand Denis, Jeff L. Froula, Zhong Wang, Robert Egan, Dongwan Don Kang, Jeffrey J. Cook, Charles Deltel, Michael Beckstette, Claire Lemaitre, Pierre Peterlongo, Guillaume Rizk, Dominique Lavenier, Yu-Wei Wu, Steven W. Singer, Chirag Jain, Marc Strous, Heiner Klingenberg, Peter Meinicke, Michael Barton, Thomas Lingner, Hsin-Hung Lin, Yu-Chieh Liao, Genivaldo Gueiros Z. Silva, Daniel A. Cuevas, Robert A. Edwards, Surya Saha, Vitor C. Piro, Bernhard Y. Renard, Mihai Pop, Hans-Peter Klenk, Markus Göker, Nikos C. Kyrpides, Tanja Woyke, Julia A. Vorholt, Paul Schulze-Lefert, Edward M. Rubin, Aaron E. Darling, Thomas Rattei, Alice C. McHardy

## Abstract

In metagenome analysis, computational methods for assembly, taxonomic profiling and binning are key components facilitating downstream biological data interpretation. However, a lack of consensus about benchmarking datasets and evaluation metrics complicates proper performance assessment. The Critical Assessment of Metagenome Interpretation (CAMI) challenge has engaged the global developer community to benchmark their programs on datasets of unprecedented complexity and realism. Benchmark metagenomes were generated from ~700 newly sequenced microorganisms and ~600 novel viruses and plasmids, including genomes with varying degrees of relatedness to each other and to publicly available ones and representing common experimental setups. Across all datasets, assembly and genome binning programs performed well for species represented by individual genomes, while performance was substantially affected by the presence of related strains. Taxonomic profiling and binning programs were proficient at high taxonomic ranks, with a notable performance decrease below the family level. Parameter settings substantially impacted performances, underscoring the importance of program reproducibility. While highlighting current challenges in computational metagenomics, the CAMI results provide a roadmap for software selection to answer specific research questions.

## Introduction

The biological interpretation of metagenomes relies on sophisticated computational analyses such as read assembly, binning and taxonomic profiling. All subsequent analyses can only be as meaningful as the outcome of these initial data processing steps. Tremendous progress has been achieved in metagenome software development in recent years^1^. However, no current approach can completely recover the complex information encoded in metagenomes. Methods often rely on simplifying assumptions that may lead to limitations and inaccuracies. A typical example is the classification of sequences into Operational Taxonomic Units (OTUs) that neglects the phenotypic and genomic diversity found within such taxonomic groupings^2^. Evaluation of computational methods in metagenomics has so far been largely limited to publications presenting novel or improved tools. However, these results are extremely difficult to compare, due to the varying evaluation strategies, benchmark datasets, and performance criteria used in different studies. Users are thus not well informed about general and specific limitations of computational methods, and their applicability to different research questions and datasets. This may result in difficulties selecting the most appropriate software for a given task, as well as misinterpretations of computational predictions. Furthermore, due to lack of regularly updated benchmarks within the community, method developers currently need to individually evaluate existing approaches to assess the value of novel algorithms or methodological improvements. Due to the extensive activity in the field, performing such evaluations represents a moving target, and consumes substantial time and computational resources, and may introduce unintended biases.

We tackle these challenges with a new community-driven initiative for the Critical Assessment of Metagenome Interpretation (CAMI). CAMI aims to evaluate computational methods for metagenome analysis comprehensively and most objectively. To enable a comprehensive performance overview, we have organized a benchmarking challenge on datasets of unprecedented complexity and degree of realism. Although comparative benchmarking has been done before^3,4^, this is the first time it has been performed as a community-driven effort. CAMI seeks to establish consensus on performance evaluation and to facilitate objective assessment of newly developed programs in the future through community involvement in the design of benchmarking datasets, evaluation procedures, choice of performance metrics, and specific questions to focus on.

We assessed the performance of metagenome assembly, binning and taxonomic profiling programs when encountering some of the major challenges commonly observed in metagenomics. For instance, the study of microbial communities benefits from the ability to recover genomes of individual strains from metagenome samples^2,5^. This enables fine-grained analyses of the functions of community members, studies of their association with phenotypes and environments, as well as understanding of the microevolution and dynamics in response to environmental changes (e.g. SNPs, lateral gene transfer, genes under directional selection, selective sweeps^6,7^ or strain displacement in fecal microbiota transplants^8^). In many ecosystems, a high degree of strain-level heterogeneity is observed^9,10^. To date, it is not clear how much assembly, genome binning and profiling software are influenced by factors such as the evolutionary relatedness of organisms present, varying community complexity, the presence of poorly categorized taxonomic groups such as viruses, or the specific parameters of the algorithms being used.

To address these questions, we generated extensive metagenome benchmarking datasets employing newly sequenced genomes of approximately 700 microbial isolates and 600 complete plasmids, viruses, and other circular elements, which were not publicly available at the time of the challenge and include organisms that are evolutionarily distinct from strains, species, genera, or orders already represented in public sequence databases. Using these genomes, benchmark datasets were designed to mimic commonly used experimental settings in the field. They include frequent properties of real datasets, such as the presence of multiple, closely related strains, of plasmid and viral sequences, and realistic abundance profiles. For reproducibility, CAMI challenge participants were encouraged to provide their predictions together with an executable docker-biobox implementing their software with specification of parameter settings and reference databases used. Overall 215 submissions representing 25 computational metagenomics programs and 36 biobox implementations of 17 participating teams from around the world were received with consent to publish. To facilitate future comparative benchmarking, all data sets are provided for download together with the current submissions in the CAMI benchmarking platform (https://data.cami-challenge.org/), allowing to submit predictions for further programs and computation of a range of performance metrics. Our results supply users and developers with extensive data about the performance of common computational methods on multiple datasets. Furthermore, we provide guidance for the application of programs, their result interpretation and suggest directions for future work.

## RESULTS

### Assembly challenge

Assembling genome sequences from short-read data remains a computational challenge, even for microbial isolates. Assembling genomes from metagenomes is even more challenging, as the number of genomes in the sample is unknown and closely related genomes occur, such as from multiple strains of the same species, potentially representing genome-sized repeats that are challenging to resolve. Nevertheless, sequence assembly is a crucial part of metagenome analysis and subsequent analyses – such as binning – depend on the quality of assembled contigs.

#### Overall performance trends

Developers submitted reproducible results for six assemblers and assembly pipelines, namely for Megahit^11^, Minia^12^, Meraga (Meraculous^13^ + Megahit), A* (using the OperaMS Scaffolder^14^), Ray Meta^15^ and Velour^16^. Several of these were specifically developed for metagenomics, while others are more broadly used (Table 1, Supplementary Table 1). The assembly results were evaluated using the MetaQUAST^17^ metrics and the reference genome and circular element sequences of the benchmark datasets (Supplementary Table 2, Supplementary methods “Assembly metrics”). As performance metrics, we focused on genome fraction and assembly size, as well as on the number of unaligned bases and misassemblies. Genome fraction measures the assembled percentage of an individual reference genome, assembly size denotes the total length in bp for an assembly (including misassembled contigs), and the number of misassemblies and unaligned bases are error metrics reflective of the assembly quality. Combined, they provide an indication of the performance of a program, while individually, they are not sufficient for assessment. For instance, while assembly size might be large, a high-quality assembly also requires the number of misassemblies and unaligned bases to be low. To assess how much metagenome data was included in each assembly, we also mapped all reads back to them.

**Table 1:**
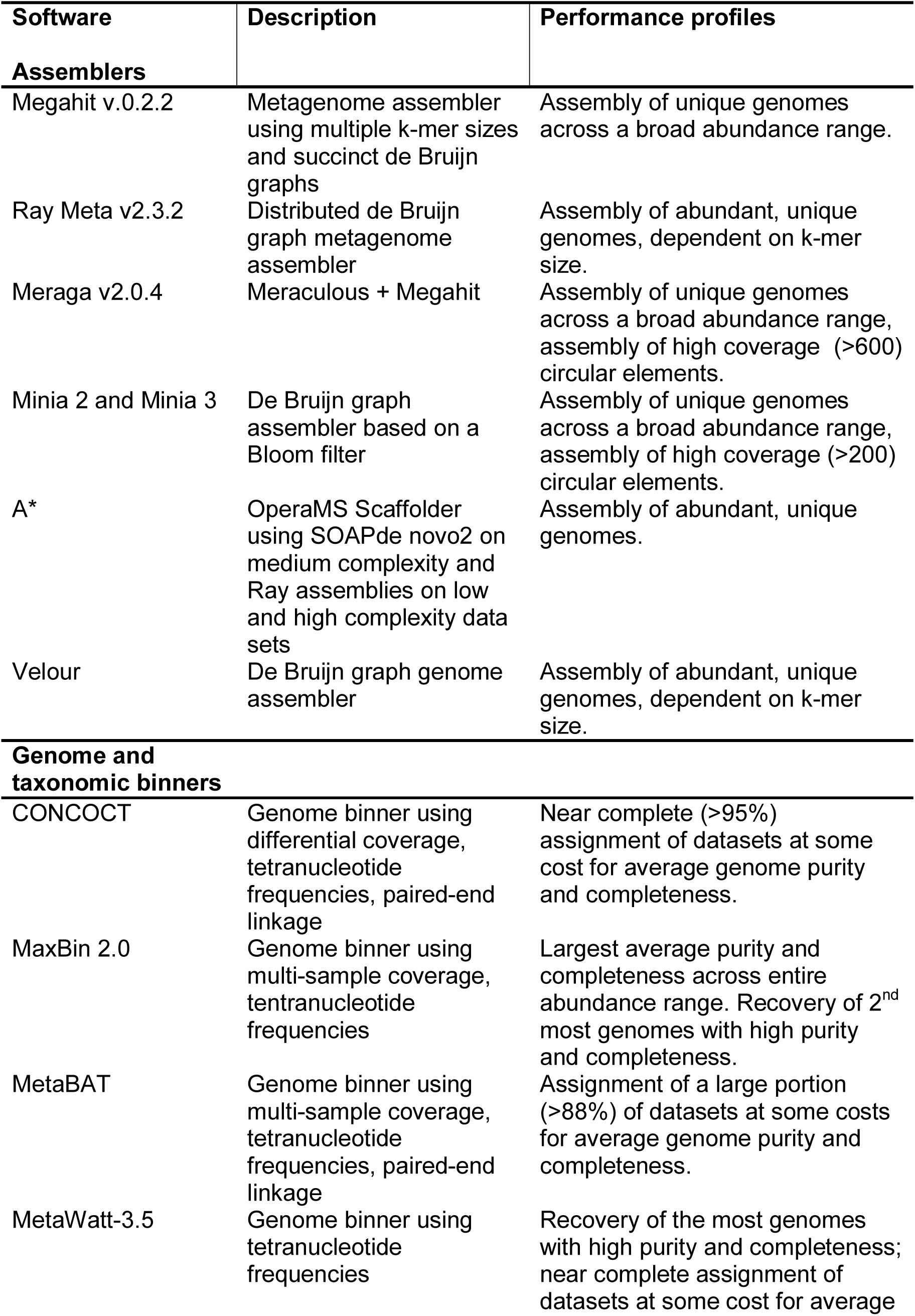
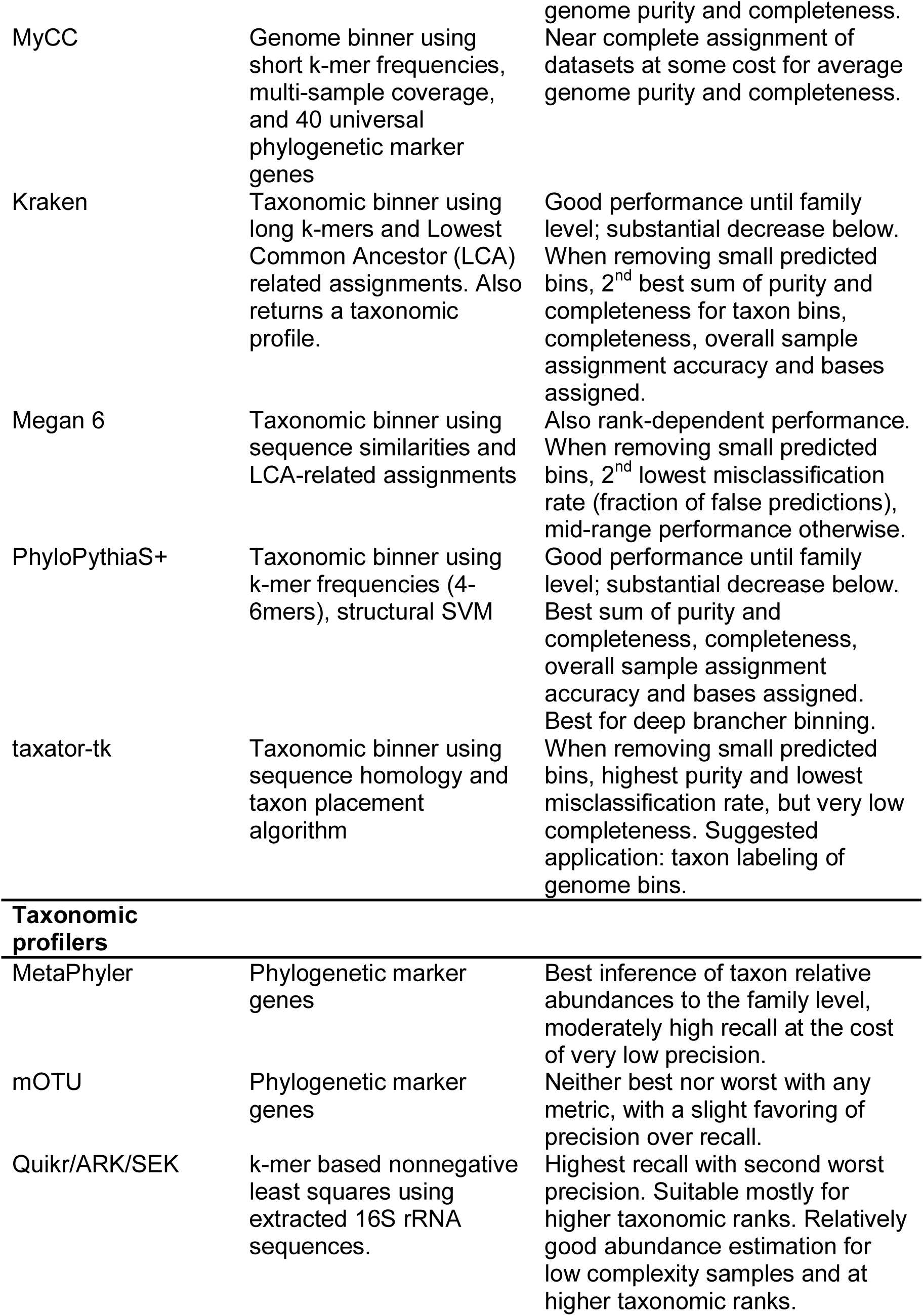
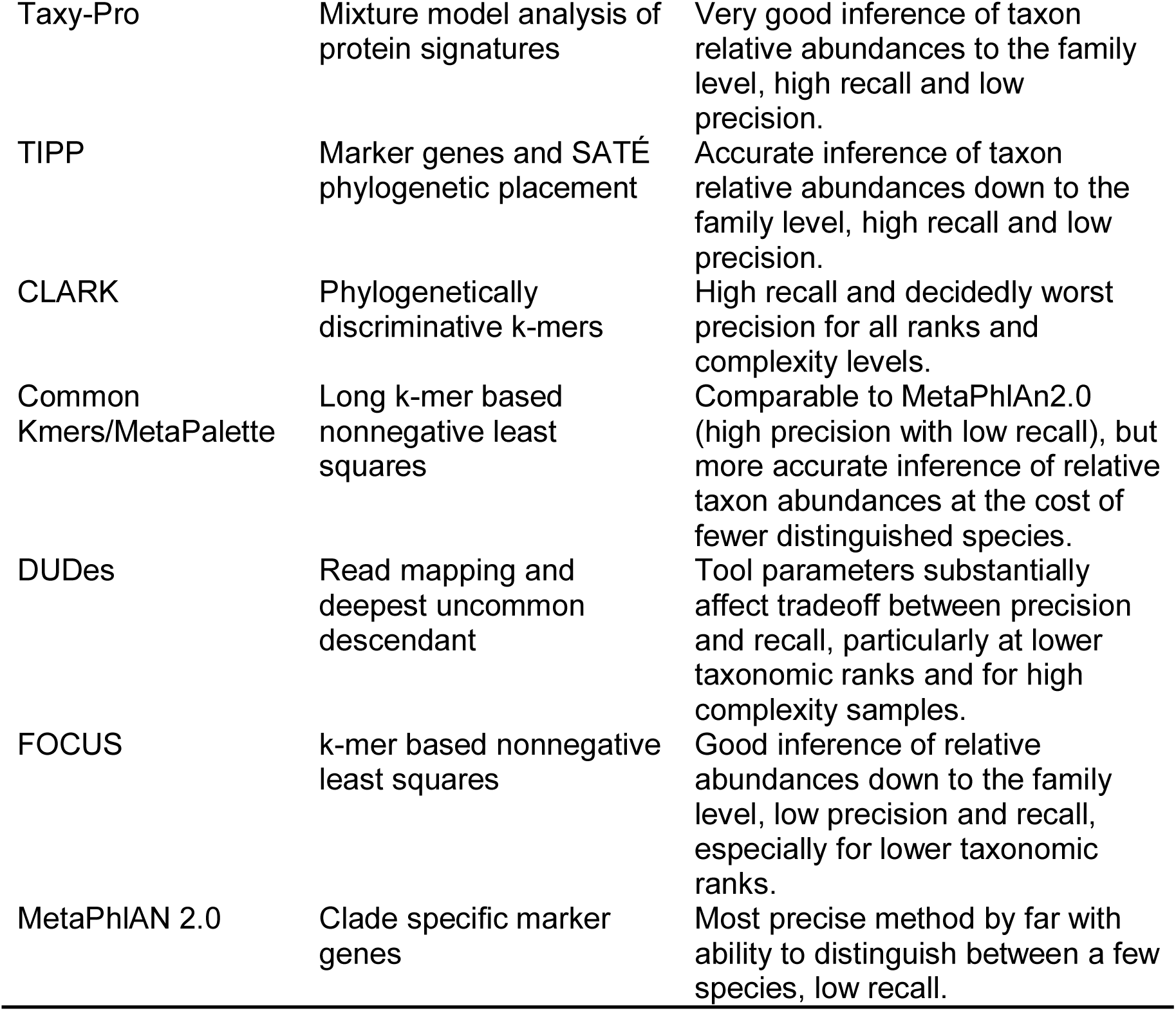
Computational metagenomics programs evaluated in the CAMI challenge. See Supplementary Tables S16 and S17 for detailed genome and taxon binning performance statistics.

Across all datasets (Supplementary Table 3) the assembly statistics varied substantially by program and parameter settings (Supplementary Figures SA1-SA12). The gold standard co-assembly of the five samples from the high complexity data set has 2.80 Gbp in 39,140 contigs. For the assemblers, values for this data set ranged from 12.32 Mbp to 1.97 Gbp assembly size (0.4% - 70% of the gold standard co-assembly, respectively), 0.4% to 69.4% genome fraction, 11 to 8,831 misassemblies and 249 bp to 40.1 Mbp unaligned contigs (Supplementary Table 2, Supplementary Fig. SA1). Megahit^11^ (*Megahit*) produced the largest assembly of 1.97 Gb, with 587,607 contigs, 69.3% genome fraction, and 96.9% mapped reads. It had a substantial number of unaligned bases (2.28 Mbp) and the largest number of misassemblies (8,831). Changing the parameters of Megahit (*Megahit_ep_mtl200*) substantially increased the unaligned bases to 40.89 Mbp, while the total assembly length, genome fraction and fraction of mapped reads remained almost identical (1.94 Gbp, 67.3%, and 97.0%, respectively, number of misassemblies: 7,538). The second largest assembly was generated by Minia^12^ (1.85 Gbp in 574,094 contigs), with a genome fraction of 65.7%, only 0.12 Mbp of unaligned bases and 1,555 misassemblies. Of all reads, 88.1% mapped to the Minia assembly. Meraga generated an assembly of 1.81 Gbp in 745,109 contigs, to which 90.5% of reads could be mapped (2.6 Mbp unaligned, 64.0% genome fraction, 2,334 misassemblies). Velour (*VELOUR_k63_C2.0*) produced the most contigs (842,405) in a 1.1 Gb assembly (15.0% genome fraction), with 381 misassemblies and 56 kbp unaligned sequences. 81% of the reads mapped to the Velour assembly. The smallest assembly was generated by Ray^6^ using *k*-mer of 91 (*Ray_k91*) with 12.3 Mbp assembled into 13,847 contigs (genome fraction <0.1%). Only 3.2% of the reads mapped to this assembly. Altogether, we found that Megahit, Minia and Meraga produced results within a similar quality range when considering these various metrics, generated a higher contiguity for the assemblies (Supplementary Figures SA10-SA12) and assembled a substantial part of the genomes across a broad range of abundances. Analysis of the low and medium complexity data sets delivered similar results (Supplementary Figs SA4-SA6, SA7-SA9).

#### Closely related genomes

To assess how the presence of closely related genomes affects the performance of assembly programs, we divided genomes according to their Average Nucleotide Identity (ANI^18^) to each other into “unique strains” (genomes with < 95% ANI to any other genome) and “common strains” (genomes with closely related strains present; all genomes with an ANI >= 95% to any other genome in the dataset). When considering the fraction of all reference genomes recovered, Meraga, Megahit and Minia performed best (Fig. 1**a**). For the unique strains, Minia and Megahit had the highest genome recovery rate (Fig. 1**c**; median over all genomes 98.2%), followed by Meraga (median 96%) and *VELOUR_k31_C2.0* (median 62.9%). Notably, for the common strains, the recovery rate dropped substantially for all assemblers (Fig. 1**b**). Megahit (*Megahit_ep_mtl200*) recovered this group of genomes best (median 22.5%), followed by Meraga (median 12.0%) and Minia (median 11.6%). *VELOUR_k31_C2.0* showed only a genome fraction of 4.1% (median) for this group of genomes. Thus, current metagenome assemblers produce high quality results for genomes for which no close relatives are present. Only a small fraction of the “common strain” genomes was assembled, with assembler-specific differences. For very high ANI groups (>99.9%), most assemblers recovered individual genomes (Supplementary Fig. SA16). The resolution of strain-level diversity represents a substantial challenge to all evaluated programs.

**Figure 1:**
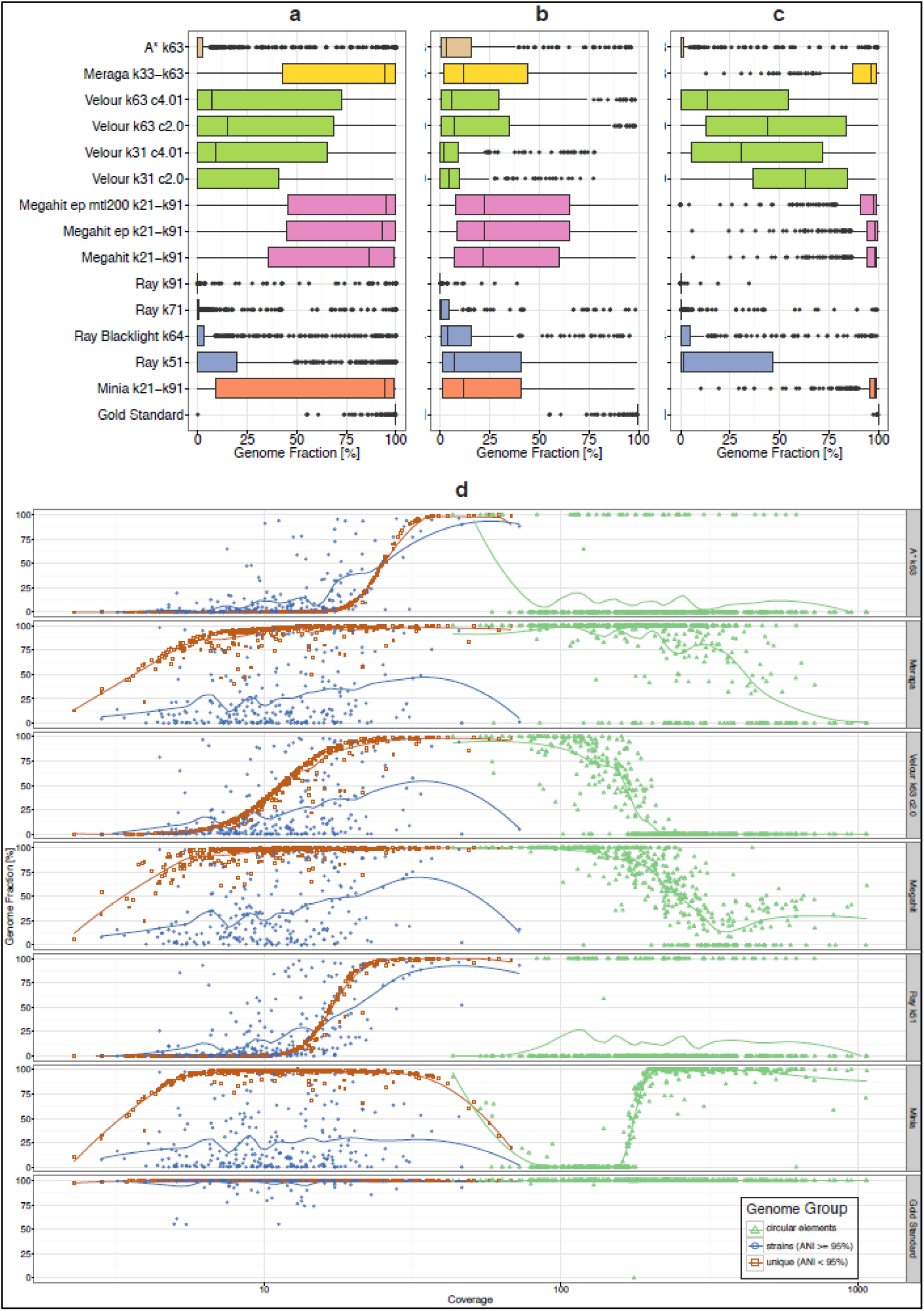
Boxplots representing the fraction of reference genomes assembled by each assembler for the high complexity data set. (**a**): all genomes, (**b**): genomes with ANI >=95%, (**c**): genomes with ANI < 95%. Coloring indicates the results from the same assembler incorporated in different pipelines or with other parameter settings. (**d**): genome recovery fraction versus genome sequencing depth (coverage) for the high complexity data set. Data were classified as unique genomes (ANI < 95%, brown color), genomes with related strains present (ANI >= 95%, blue color) and high copy circular elements (green color). The gold standard includes all genomic regions covered by at least one read in the metagenome dataset, therefore the genome fraction for low abundance genomes can be less than 100%.

#### Effect of sequencing depth

To investigate the effect of sequencing depth on the assembly metrics, we compared the genome recovery rate (genome fraction) to the genome sequencing coverage for the gold standard and all assemblies (Fig. 1**d**, Supplementary Fig. SA2 for complete results). The chosen *k*-mer size affects the recovery rate (Supplementary Fig. SA3): while small *k*-mers allowed an improved recovery of low abundance genomes, large *k*-mers led to a better recovery of highly abundant ones. Assemblers using multiple *k*-mers (Minia, Megahit, Meraga) substantially outperformed single *k*-mer assemblers. Most assemblers poorly recovered very high copy circular elements (sequencing coverage > 100x), except for Meraga and the Minia Pipeline, which both performed well for a substantial portion, though Minia surprisingly lost all genomes with a sequencing coverage between 80 and 200x (Fig. 1**d**). Notably, no program investigated contig topology, and determined whether these were circular and complete.

### Binning challenge

Metagenome assembly programs return mixtures of variable length fragments originating from individual genomes. Metagenome binning algorithms were devised to tackle the problem of classifying, or “binning” these fragments according to their genomic or taxonomic origins. These “bins”, or sets of assembled sequences and reads, group data from the genomes of individual strains or of higher-ranking taxa present in the sequenced microbial community. Such bin reconstruction allows the subsequent analysis of the genomes (or pangenomes) of a strain (or higher-ranking taxon) from a microbial community. While genome binners group sequences into unlabeled genome bins, taxonomic binners group the sequences into bins with a taxonomic label attached.

Results for five genome binners and four taxonomic binners were submitted together with bioboxes of the respective programs in the CAMI challenge, namely for MyCC^19^, MaxBin 2.0^20^, MetaBAT^21^, MetaWatt-3.5^22^, CONCOCT^23^, PhyloPythiaS+^24^, taxator-tk^25^, MEGAN 6^26^ and Kraken^27^. Submitters could choose to run their program on the provided gold standard assemblies or on individual read samples (MEGAN 6), according to their suggested application. We then determined their performance for addressing important questions in microbial community studies: do they allow the recovery of high quality genome bins for individual strains, i.e. with high average completeness (recall), and purity (precision), i.e. low contamination levels? How does strain level diversity affect performance? How is performance affected by the presence of non-bacterial sequences in a sample, such as viruses or plasmids? Do current taxonomic binners allow recovery of higher-ranking taxon bins with high quality? How does their performance vary across taxonomic ranks? Which programs are highly precise in taxonomic assignment, so that their outputs can be used to assign taxa to genome bins? Which software has high recall in the detection of taxon bins from low abundance community members, as is required for metagenomes from ancient DNA and for pathogen detection? Finally, which programs perform well in the recovery of bins from deep-branching taxa, for which no sequenced genomes yet exist?

#### Recovery of individual genome bins

We first investigated the performance of each program in the recovery of individual genome (strain-level) bins. We calculated completeness and purity (Supplementary Methods) for every bin relative to the genome that was most abundant in that bin in terms of assigned sequence length. In addition, we calculated the Adjusted Rand Index (ARI) as measure of assignment accuracy for the portion of the data assigned by the different programs. As not all programs assigned the entire data set to genome bins, these values should be interpreted under consideration of the fraction of data assigned (Fig. 2**d**). These two measures complement completeness and purity averaged over genome bins, as assignment accuracy is evaluated per bp, with large bins contributing more than smaller bins in the evaluation.

**Figure 2:**
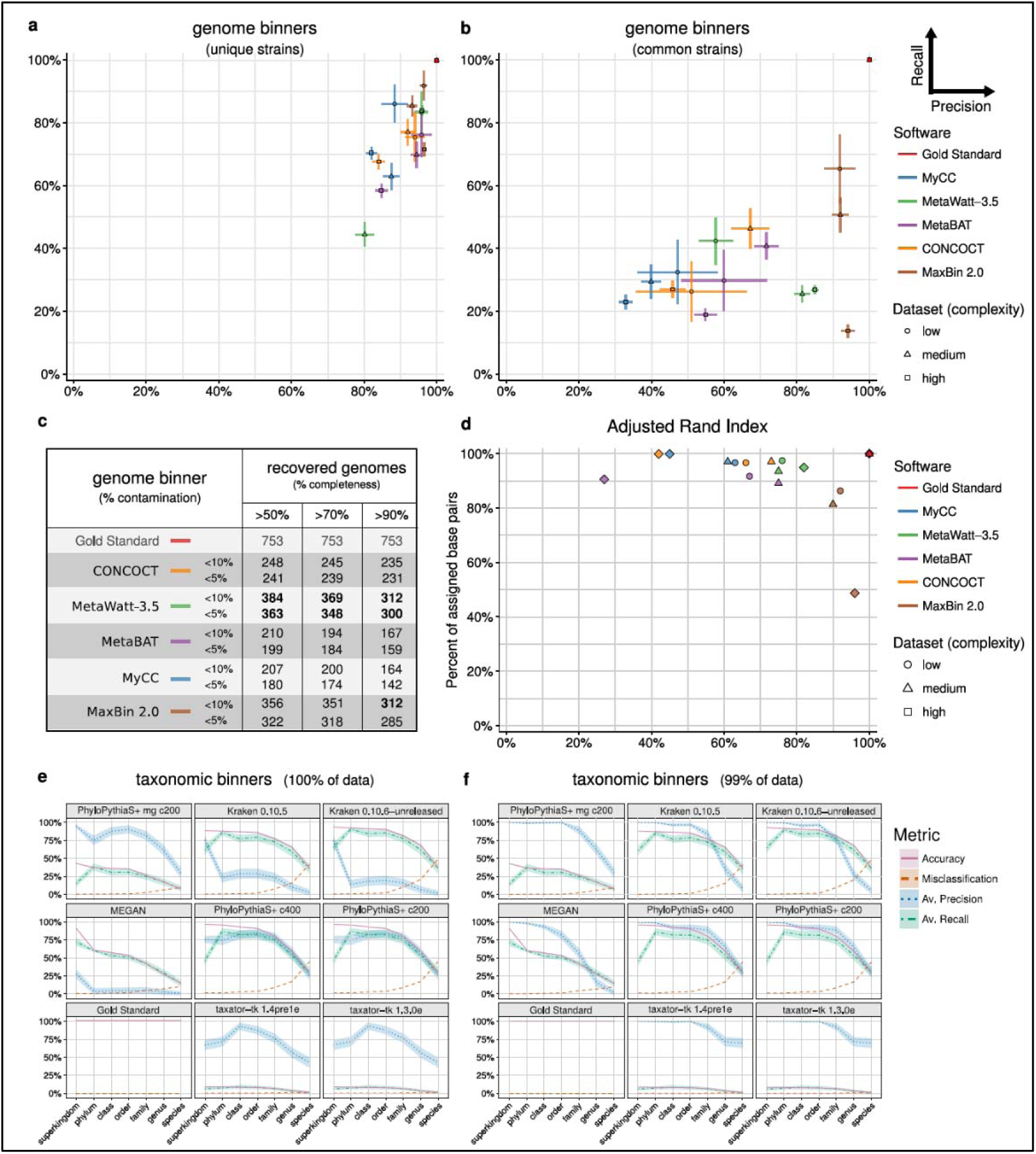
Average purity (x-axis) and completeness (y-axis) and their standard errors (bars) for genomes reconstructed by genome binners; for genomes of unique strains with equal to or less than 95% ANI to others (**a**) and common strains with more than 90% ANI to each other (**b**). For each program and complexity dataset, the submission with the largest sum of purity and completeness is shown (Supplementary Tables 1, 10, 12). In each case, small bins adding up to 1% of the data set size were removed. (**c**) Number of genomes recovered with varying completeness and contamination (1-purity, Supplementary Table S17). (**d**) The Adjusted Rand Index (ARI, x-axis) in relation to fraction of the sample assigned (in basepairs) by the genome binners (y-axis). The ARI was calculated excluding unassigned sequences, thus reflects the assignment accuracy for the portion of the data assigned. (**e**,**f**) Taxonomic binning performance metrics across ranks for the medium complexity data set, with (**e**) results for the complete data set and (**f**) with smallest predicted bins summing up to 1% of the data set removed. Shaded areas indicate the standard error of the mean in precision (purity) and recall (completeness) across taxon bins.

For the genome binners, both average bin completeness (ranging from 34% to 80%) and purity (ranging from 70% to 97%) varied substantially across the three datasets (Supplementary Table 4, Supplementary Fig. B1). For the medium and low complexity datasets, MaxBin 2.0 had the highest average values (70-80% completeness, more than >92% purity), followed by other programs with comparably good performance in a narrow range (completeness ranging with one exception from 50-64%, more than 75% purity). Notably, other programs assigned a larger portion of the datasets in bp than MaxBin 2.0, though with lower ARI (Fig. 2**d**). For applications where binning a larger fraction of the dataset at the cost of some accuracy is important, therefore, programs such as MetaWatt-3.5, MetaBAT and CONCOCT could be a good choice. The high complexity dataset was more challenging to all programs, with average genome completeness decreasing to around 50% and more than 70% purity, except for MaxBin 2.0 and MetaWatt-3.5, which showed an outstanding purity of above 90%. The programs either assigned only a smaller dataset portion (>50% of the sample bp, MaxBin 2.0) with high ARI or assigned a larger fraction with lower ARI (more than 90% with less than 0.5 ARI). The exception was MetaWatt-3.5, which assigned more than 90% of the dataset with an ARI larger than 0.8, thus best recovering abundant genomes from the high complexity dataset. Overall, MetaWatt-3.5, closely followed by MaxBin 2.0, recovered the most genomes with high purity and completeness from the three datasets (Fig. 2**c**, Supplementary Table 17).

#### Effect of strain diversity

When considering only unique strains, the performance of all genome binners improved substantially, both in terms of average purity and completeness per genome bin (Fig. 2**a**). For the medium and low complexity datasets, all had a purity of above 80%, while completeness was more variable. MaxBin 2.0 performed best across all three datasets, showing more than 90% purity and 70% or more completeness. An almost equally good performance for two datasets was delivered by MetaBAT, CONCOCT and MetaWatt-3.5.

For the "common strains" of all three datasets, however, completeness decreased substantially (Fig. 2**b**), similarly to purity for most programs. MaxBin 2.0 still stood out, with more than 90% purity on all datasets. Interestingly, when considering the value of bins reconstructed by taxon binners for genome reconstruction, taxonomic binners had lower completeness than genome binners, but reached a similar purity, thus delivering high quality, partial genome bins (Supplementary Material 1.4.4; Supplementary Fig. B9). Overall, the presence of multiple related strains in a metagenome sample had a substantial effect on the quality of the reconstructed genome bins, both for genome and taxonomic binners. Very high quality genome bin reconstructions were attainable with genome binning programs for “unique” strains, while the presence of several closely related strains presented a notable hurdle to these tools.

#### Performance in taxonomic binning

We next investigated the performance of taxonomic binners in recovering taxon bins at different ranks. These results can be used for taxon-level evolutionary or functional pangenome analyses and conversion into taxonomic profiles. As performance metrics, the average purity (precision) and completeness (recall) per taxon bin were calculated for individual ranks under consideration of the taxon assignment (Supplementary Material, Binning metrics). In addition, we determined the overall classification accuracy for each dataset, as measured by total assigned sequence length, and misclassification rate for all assignments. While the former two measures allow assessing performance averaged over bins, where all bins are treated equally, irrespective of their size, the latter are influenced by the actual sample taxonomic constitution, with large bins having a proportionally larger influence.

For the low complexity data set, PhyloPythiaS+ had the highest sample assignment accuracy, average taxon bin completeness and purity, which were all above 75% from domain to family level. Kraken followed, with average completeness and accuracy still above 50% until family level. However, purity was notably lower, mostly caused by prediction of many small false bins, which affects purity more than overall accuracy, as explained above (Supplementary Fig. B3). Removing the smallest predicted bins (1% of the data set) increased purity for Kraken, MEGAN, and, most strongly, for taxator-tk, for which it was close to 100% until the order level, and above 75% until the family level (Supplementary Fig. B4). This shows that small predicted bins by these programs are not reliable, but otherwise, high purity can be reached for higher ranks. Below the family level no program performed very well, with all either assigning very little data (low completeness and accuracy, accompanied by a low misclassification rate), or performing more assignments with a substantial amount of misclassification. Another interesting observation is the similar performance for Kraken and Megan, which was not observed on the other datasets, though. These programs employ different features of the data (Table 1), but rely on similar algorithms.

The results for the medium complexity data set qualitatively agreed with those obtained for the low complexity data set, except that Kraken, MEGAN and taxator-tk performed better (Fig. 2**e**). With the smallest predicted bins removed, both Kraken and PhyloPythiaS+ performed similarly well, reaching above 75% for accuracy, average completeness and purity until the family rank (Fig. 2**f**). Similarly, taxator-tk showed an average purity of almost 75% even down to the genus level (almost 100% until order level) and MEGAN more than 75% down to the order level, while maintaining accuracy and average completeness of around 50%. The results of high purity taxonomic predictions can be combined with genome bins, to enable their taxonomic labeling. The performance for the high complexity data set was similar to that for the medium complexity data set (Supplementary Figs. B5, B6).

#### Analysis of low abundance taxa

We determined which programs had high completeness also for low abundance taxa. This is relevant when screening for pathogens in diagnostic settings^28^, or for metagenome studies of ancient DNA samples. Even though a high completeness was achieved by PhyloPythiaS+ and Kraken until the rank of family (Fig. 2**e**,**f**), it degraded for lower ranks and low abundance bins (Supplementary Fig. B7), which are of most interest for these applications. It therefore remains a challenge to further improve the predictive performance.

#### Deep-branchers

Taxonomic binning methods commonly rely on comparisons to reference sequences for taxonomic assignment. To investigate the effect of increasing evolutionary distances between a query sequence and available genomes, we partitioned the challenge datasets by their taxonomic distances to sequenced reference genomes as genomes of new strains, species, genus or family (Supplementary Fig. B8). For genomes representing new strains from sequenced species, all programs performed well, with generally high purity and oftentimes high completeness, or with characteristics observed also for other datasets (such as low completeness for taxator-tk). At increasing taxonomic distances to the reference, performance for MEGAN and Kraken dropped substantially, in terms of both purity and completeness, while PhyloPythiaS+ decreased most notably in purity and taxator-tk in completeness. For deep branchers at larger taxonomic distances to the reference collections, PhyloPythiaS+ maintained the best overall purity and completeness.

#### Influence of plasmids and viruses

The presence of plasmid and viral sequences had almost no effect on the performance for binning bacterial and archaeal organisms. Although the copy numbers of plasmids and viral data were high, in terms of sequence size, the fraction of viral, plasmid and other circular elements was small (<1.5%, Supplementary Table 6). Only Kraken and MEGAN 6 made predictions for the viral fraction of the data or predicted viruses to be present, though with low purity (<30%) and completeness (<20%).

### Profiling challenge

Taxonomic profilers predict the identity and relative abundance of the organisms (or higher level taxa) from a microbial community using a metagenome sample. This does not result in classification labels for individual reads or contigs, which is the aim of taxonomic binning methods. Instead, taxonomic profiling is used to study the composition, diversity, and dynamics of clusters of distinct communities of organisms in a variety of environments^29-31^. In some use cases, such as identification of potentially pathogenic organisms, accurate determination of the presence or absence of a particular taxon is important. In comparative studies (such as quantifying the dynamics of a microbial community over an ecological gradient), accurately determining the relative abundance of organisms is paramount.

Members of the community submitted results for ten taxonomic profilers to the CAMI challenge: CLARK^32^; ‘Common kmers’ (an early version of MetaPalette^33^, abbreviated CK in the figures); DUDes^34^; FOCUS^35^; MetaPhlAn 2.0^36^; Metaphyler^37^; mOTU^38^; a combination of Quikr^39^, ARK^40^, and SEK^41^ (abbreviated Quikr); Taxy-Pro^42^; and TIPP^43^. For several programs, results were submitted with multiple versions or different parameter settings, bringing the number of unique submissions to twenty.

#### Performance trends

We employed commonly used metrics (Supplementary Material ‘Profiling Metrics’) to assess the quality of taxonomic profiling submissions with regard to the biological questions outlined above. These can be divided into abundance metrics (L1 norm and weighted Unifrac^44^) and binary classification measures (true positives, false positives, false negatives, recall, and precision). In short, the abundance metrics assess how well a particular method reconstructs the relative abundances in comparison to the gold standard, with the L1 norm using the sum of differences in abundances (ranges between 0 and 2) and Unifrac using differences weighted by distance in the taxonomic tree (ranges between 0 and 16). The binary classification metrics assess how well a particular method detects the presence or absence of an organism in comparison to the gold standard, irrespective of their abundances. All metrics except the Unifrac metric (which is rank independent) are defined at each taxonomic rank.

We observed a large degree of variability in reconstruction fidelity for all profilers across metrics, taxonomic ranks, and samples. Each had a unique error profile, with different profilers showing different strengths and weaknesses (Fig. 3**a**). In spite of this variability, when comparing results for each sample, a number of patterns emerged. The profilers could be placed in three categories: (1) profilers that correctly predicted the relative abundances, (2) precise ones, and (3) profilers with high recall. To quantify this observation, we determined the following summary statistics: for each metric, on each sample, we ranked the profilers by their performance. Each was assigned a score for its ranking (0 for first place among all tools at a particular taxonomic rank for a particular sample, 1 for second place, etc.). These scores were then added over the taxonomic ranks to the genus level and summed over the samples, to give a global performance score (Fig. 3**b**, Supplementary Figs P1-P7, Supplementary Table 7).

**Figure 3:**
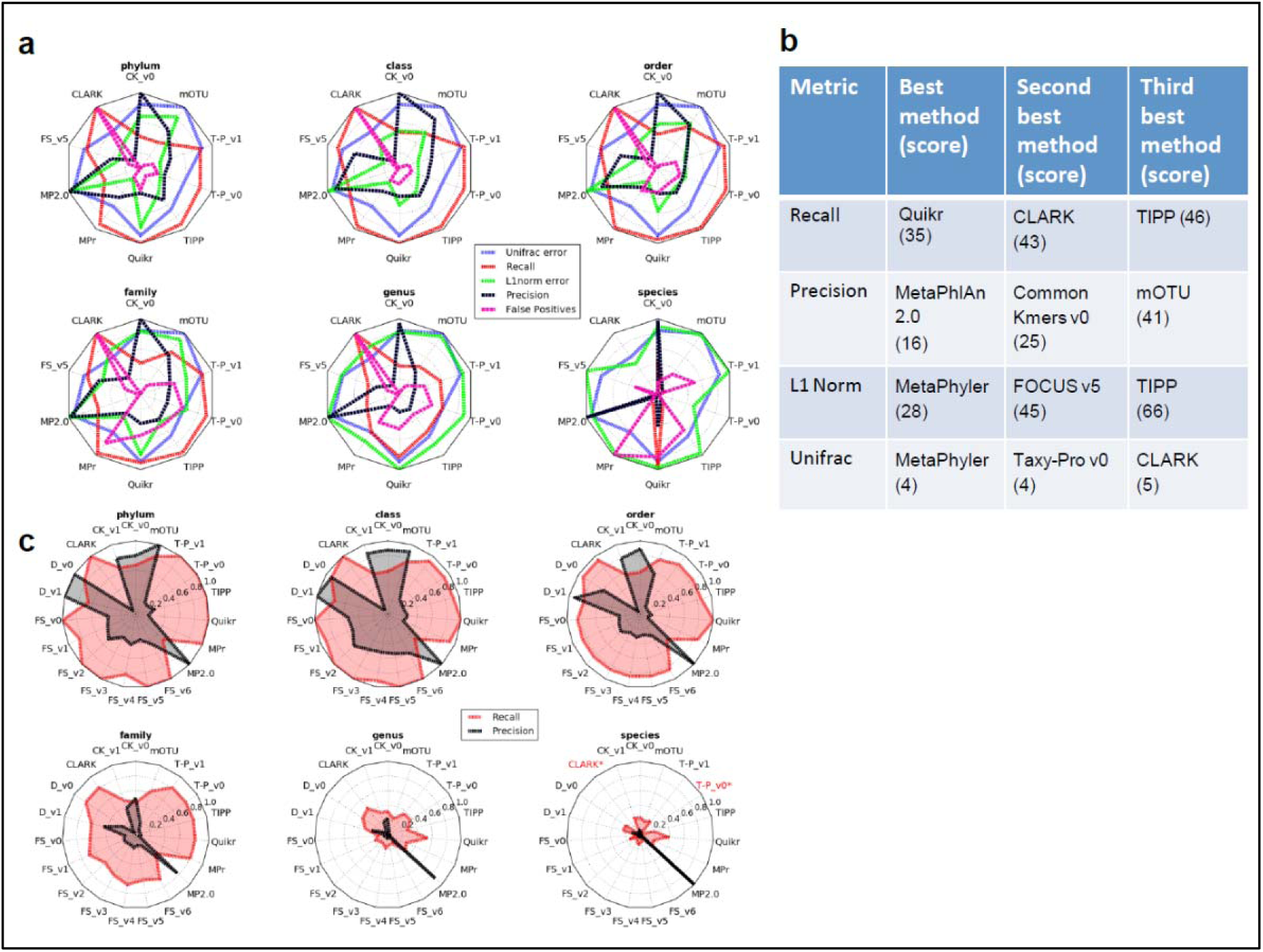
(**a**) Relative performance of profilers for different ranks and with different error metrics (weighted Unifrac, L1 norm, recall, precision, and false positives), shown here exemplarily for the microbial portion of the first high complexity sample. Each error metric was divided by its maximal value to facilitate viewing on the same scale and relative performance comparisons. A method’s name is given in red (with two asterisks) if it returned no predictions at the corresponding taxonomic rank. (**b**) Best scoring profilers using different performance metrics summed over all samples and taxonomic ranks to the genus level. A lower score indicates that a method was more frequently ranked highly for a particular metric. The maximum (worst) score for the Unifrac metric is 38 = (18 + 11 + 9) profiling submissions for the low, medium and high complexity datasets respectively), while the maximum score is 190 for all other metrics (= 5 taxonomic ranks * (18 + 11 + 9) profiling submissions for the low, medium and high complexity datasets respectively). (**c**) Absolute recall and precision for each profiler on the microbial (filtered) portion of the low complexity data set across six taxonomic ranks. Abbreviations are FS (FOCUS), T-P (Taxy-Pro), MP2.0 (MetaPhlAn 2.0), MPr (Metaphyler), CK (Common Kmers) and D (DUDes).

The profilers with the highest recall were Quikr, CLARK, Tipp, and Taxy-Pro (Fig. 3), indicating their suitability for pathogen detection, where failure to identify an organism can have severe negative consequences. The profilers with the highest recall were also among the least precise (Supplementary Figs P8-P12) where low precision was typically due to prediction of a large number of low abundance organisms. In terms of precision, MetaPhlAn 2.0 and “Common Kmers” demonstrated an overall superior performance, indicating that these two are best at only predicting organisms that are actually present in a given sample and suggesting their use in scenarios where many false positives can cause unwanted increases in costs and effort in downstream analysis. The programs that best reconstructed the relative abundances were MetaPhyler, FOCUS, TIPP, Taxy-Pro, and CLARK, making such profilers desirable for analyzing organismal abundances between and among metagenomic samples.

Often, a balance between precision and recall is desired. To assess this, we took for each profiler one half of the sum of precision and recall and averaged this over all samples and taxonomic ranks. The top performing programs by this criterion were Taxy-Pro v0, (mean=0.616), MetaPhlAn 2.0 (mean=0.603), and DUDes v0 (mean=0.596).

#### Performance at different taxonomic ranks

Most profilers performed well at higher taxonomic ranks (Fig. 3**c** and Supplementary Figs. P8-P12). A high recall was achieved until family level, and degraded substantially below. For example, over all samples and tools at the phylum level, the mean±SD recall was 0.845±0.194, and the median L1 norm was 0.382±0.280, both values close to each of these metrics’ optimal value (ranging from 1 to 0 and 0 to 2, respectively). The precision had the largest variability among the metrics, with a mean phylum level value of 0.529 with a standard deviation of 0.549. Precision and recall were simultaneously high for several methods (DUDes, Common Kmers, mOTU, and MetaPhlAn 2.0) until the rank of order. We observed that accurately reconstructing a taxonomic profile is still difficult for the genus level and below. Even for the low complexity sample, only MetaPhlAn 2.0 maintained its precision down to the species level, while the maximum recall at genus rank for the low complexity sample was 0.545 for Quikr. Across all profilers and samples, there was a drastic average decrease in performance between the family and genus level of 47.5±14.9% and 51.6±18.1% for recall and precision, respectively. In comparison, there was little change between the order and family levels, with a decrease of only 9.7±6.9% and 8.8±26.4% for recall and precision, respectively. The other error metrics showed similar performance trends for all samples and methods (Figs 3**c** and Supplementary Figs. P13-P17).

#### Parameter settings and software versions

Several profilers were submitted with different parameter settings or versions (Supplementary Table 1). For some, this had little effect: for instance, the variance in recall among 7 different versions of FOCUS on the low complexity sample at the family level was only 0.002. For others, this caused large changes in performance: for instance, one version of DUDes had twice the recall compared to another at the phylum level on the pooled high complexity sample (Supplementary Figs. P13-P17). Interestingly, a few developers chose not to submit results beyond a fixed taxonomic rank, such as for Taxy-Pro and Quikr. These submissions generally performed better than default program versions submitted by the CAMI team; indicating that, not surprisingly, experts can generate better results than when using a program’s default setting.

#### Performance for viruses and plasmids

In addition to microbial sequence material, the challenge datasets also included sequences of plasmids, viruses and other circular elements (Supplementary Table 7). We investigated the effect of including these data in the gold standard profile for the taxonomic profilers (Supplementary Figs P18-P20). Here, the term “filtered” is used to indicate the gold standard did not include these data, and the term “unfiltered” indicates use of these data. The metrics affected by the presence of these data were the abundance-based metrics (L1 norm at the superkingdom level and Unifrac), and precision and recall (at the superkingdom level). All methods correctly detected Bacteria and Archaea, indicated by a recall of 1.0 at the superkingdom level on the filtered samples. The only methods to detect viruses in the unfiltered samples were MetaPhlAn 2.0 and CLARK. Averaging over all methods and samples, the L1 norm at the superkingdom level increased from 0.051 for the filtered samples to 0.287 for the unfiltered samples. Similarly, the Unifrac metric increased from 7.213 for the filtered to 12.361 for the unfiltered datasets. Thus, a substantial decrease in the fidelity of abundance estimates was caused by the presence of viruses and plasmids in a sample.

#### Taxonomic profilers vs. profiles derived from taxonomic binning

We compared the profiling results to those generated by several taxonomic binners using a simple coverage-approximation conversion algorithm for deriving profiles from taxonomic bins (Supplementary Methods, Figs P21-P24). Overall, the taxonomic binners were comparable to the profilers in terms of precision and recall. At the order level, the mean precision over all taxonomic binners was 0.595 (versus 0.401 for the profilers) and the mean recall was 0.816 (versus 0.857 for the profilers). Two binners, MEGAN 6 and PhyloPythiaS+, had better recall than the profilers at the family level, with the degradation in performance past the family level being evident for the binners as well. However for precision at the family level, PhyloPythiaS+ was the fourth, after the profilers CK_v0, MetaPhlan 2.0, and the binner taxator-tk (Supplementary Figs P21-P22).

Abundance estimation at higher ranks was more problematic for the binners, as the L1 norm error at the order level was 1.07 when averaged over all samples, while the profilers average was only 0.681. Overall, though, the binners delivered slightly more accurate abundance estimates, as the binning average Unifrac metric was 7.03, while the profiling average was 7.23. These performance differences may in part be due to the use of the gold standard contigs as input by the binners except for MEGAN 6, though oftentimes Kraken is also applied to raw reads, while the profilers used the raw reads.

## CONCLUSIONS

Determination of program performance is essential for assessing the state of the art in computational metagenomics. However, a lack of consensus about benchmarking datasets and evaluation metrics has complicated comparisons and their interpretation. To tackle this problem, CAMI has engaged the global developer community in a benchmarking challenge, with more than 40 teams initially registering for the challenge and 19 teams handing in submissions for the three different challenge parts. This was achieved by providing benchmark datasets of unprecedented complexity and degree of realism, generated exclusively from around 700 newly sequenced microbial genomes and 600 novel viruses, plasmids and other circular elements. These spanned a range of evolutionary divergences from each other and from publicly available reference collections. We implemented commonly used metrics in close collaboration with the computational and applied metagenomics communities and agreed on the metrics most important for common research questions and biological use cases in microbiome research using metagenomics. To be of practical value to researchers, the program submissions have to be reproducible, which requires knowledge of reference data, parameter settings and program versions. In CAMI, we have taken steps to ensure reproducibility by development of docker-based bioboxes^45^ and encouraging developer submissions of bioboxes for the benchmarked metagenome analysis tools, enabling their standardized execution and format usages. The benchmark datasets, along with the CAMI benchmarking platform allow further result submissions and their evaluation on the challenge data sets, to facilitate benchmarking of further programs. Currently, we are extending the platform capabilities for automated benchmarking of bioboxpackaged programs on these and further data sets, as well as comparative result visualizations.

The evaluation of assembly programs revealed a clear advantage for assemblers using a range of *k*-mers compared to single *k*-mer assemblies (Table 1). While single *k*-mer assemblies reconstructed only genomes with a certain coverage (small *k*-mers for low abundant genomes, large *k*-mers for high abundant genomes), using multiple *k*-mers significantly improved the fraction of recovered genomes. Two programs performed well in the reconstruction of high copy circular elements, although none detected their circularities. An unsolved challenge of metagenomic assembly for all programs is the reconstruction of closely related genomes. A poor assembly quality or lack of assembly for these genomes will negatively impact subsequent contig binning, as the contigs will be missing in the assembly output, further complicating their study.

In evaluation of the genome and taxonomic binners, all programs were found to perform surprisingly well at genome reconstruction, if no closely related strains were present. Taxonomic binners performed acceptably in taxon bin reconstruction down to the family rank (Table 1). This leaves a gap in species and genus-level reconstruction that is to be closed, also for taxa represented by single strains in a microbial community. Taxonomic binners achieved a better precision in genome reconstruction than in species or genus-level binning, raising the possibility that a part of the decrease of performance in low ranking taxon assignment is due to limitations of the reference taxonomy used. A sequence-derived reference phylogeny might represent a more suitable framework for – in that case – “phylogenetic” binning. When comparing the average performance of taxon binners for taxa with similar surroundings in the SILVA and NCBI reference taxonomies to those with less agreement, we observed a significant difference for taxa with discrepant surroundings, primarily a decreased performance for lower taxonomic ranks until family level (Supplementary Methods, Section 1.4.5; Supplementary Table S24). Thus, the use of SILVA might further improve taxon binning, though the lack of associated genome sequence data represents a practical hurdle^46^. Another challenge for all programs is the deconvolution of strain-level diversity, which we found to be substantially less effective than binning of genomes without close relatives present. For the typically covariance of read coverage based genome binners it may require substantially larger numbers of replicate samples than those analyzed here (up to 5) to attain a satisfactory performance.

Despite of a large variability in performance amongst the submitted profilers, most profilers performed well with good recall and low errors in abundance estimates until the family rank, with precision being the most variable of these metrics (Table 1). The use of different classification algorithms, reference taxonomies, reference databases and information sources (marker gene versus genome wide k-mer based) are likely contributors to the observed performance differences. To enable more systematic analyses of their individual impacts, software developers could provide configurable options for use of databases, k-mer sizes or other specialized settings, instead of having these hard-coded. Similarly to taxonomic binners, performance across all metrics substantially decreased for the genus level and below. Also when taking plasmids and viruses into consideration for abundances estimates, the performance of all programs decreased substantially, indicating a need for further development to enable a better analysis of datasets with such content, as plasmids are likely to be present and viral particles are not always removed by size filtration^47^.

As both the sequencing technologies and the computational metagenomics programs continue to evolve rapidly, CAMI will provide further benchmarking challenges to the community. Long read technologies such as those by Oxford Nanopore, Illumina and PacBio^48^ are expected to become more common in metagenomics, which will in turn require other assembly methods and may allow a better resolution of closely related genomes from metagenomes. In the future, we also plan to tackle assessment of runtimes and RAM requirements, to determine program suitability for different use cases, such as execution on individual desktop machines or as part of computational metagenome pipelines provided by MG-RAST^49^, EMG^50^ or IMG/M^51^. We invite everyone interested to join and work with CAMI on providing comprehensive performance overviews of the computational metagenomics toolkit, to inform developers about current challenges in computational metagenomics and applied scientists of the most suitable software for their research questions.

## ONLINE METHODS

### Community involvement

We organized public workshops, roundtables, hackathons and a research programme around CAMI at the Isaac Newton Institute for Mathematical Sciences (Supplementary Fig. M1), to decide on the principles realized in data set and challenge design. To determine the most relevant metrics for performance evaluation, a meeting with developers of evaluation software and of commonly used binning, profiling and assembly software was organized. Subsequently we created biobox containers implementing a range of commonly used performance metrics, including the ones decided as most relevant in this meeting (Supplementary Table 8). Computational support for challenge participants was provided by the Pittsburgh Supercomputing Center.

### Standardization and reproducibility

For performance assessment, we developed several standards: we defined output formats for profiling and binning tools, for which no widely accepted standard existed. Secondly, standards for submitting the software itself, along with parameter settings and required databases were defined and implemented in docker container templates named bioboxes^45^. These enable the standardized and reproducible execution of submitted programs from a particular category. Challenge participants were encouraged to submit the results together with their software in a docker container following the bioboxes standard. In addition to 23 bioboxes submitted by challenge participants, we generated 13 additional bioboxes and ran them on the challenge datasets (Supplementary Table 1), working with the developers to define the most suitable execution settings, if possible. For several submitted programs, bioboxes using default settings were created, to compare performance with default and expert chosen parameter settings. If required, the bioboxes can be rerun on the challenge datasets.

### Genome sequencing and assembly

Draft genomes of 310 type strain isolates were generated for the Genomic Encyclopedia of Type Strains at the DOE Joint Genome Institute (JGI) using Illumina standard shotgun libraries and the Illumina HiSeq 2000 platform. All general aspects of library construction and sequencing performed at the JGI can be found at http://www.jgi.doe.gov. Raw sequence data was passed through DUK, a filtering program developed at JGI, which removes known Illumina sequencing and library preparation artifacts [Mingkun L, Copeland A, Han J. DUK, unpublished, 2011]. The genome sequences of isolates from culture collections are available in the JGI genome portal (Supplementary Table 9). Additionally, 488 isolates from the root and rhizosphere of *Arabidopsis thaliana* were sequenced^9^. All sequenced environmental genomes were assembled using the A5 assembly pipeline (default parameters, version 20141120)^52^ and are available for download at https://data.camichallenge.org/participate). A quality control of all assembled genomes was performed based on tetranucleotide content analysis and taxonomic analyses (Supplementary Methods “Taxonomic annotation”), resulting in 689 genomes that were used for the challenge (Supplementary Table 9). Furthermore, we generated 1.7 Mb or 598 novel circular sequences of plasmids, viruses and other circular elements from multiple microbial community samples of rat caecum (Supplementary Methods, ‘Data generation’).

### Challenge datasets

We simulated three metagenome datasets of different organismal complexities and sizes by generating 150 bp paired-end reads with an Illumina HighSeq error profile from the genome sequences of 689 newly sequenced bacterial and archaeal isolates and 598 sequences of plasmids, viruses and other circular elements (Supplementary Methods “Metagenome simulation”; Supplementary Tables 3, 6; Supplementary Figs D1, D2). These datasets represent common experimental setups and specifics of microbial communities. They consist of a 15 Gb single sample dataset from a low complexity community with log normal abundance distribution (40 genomes and 20 circular elements), a 40 Gb differential log normal abundance dataset with two samples of a medium complexity community (132 genomes and 100 circular elements) and long and short insert sizes, as well as a 75 Gb time series dataset with five samples from a high complexity community with correlated log normal abundance distributions (596 genomes and 478 circular elements). Some important properties of the benchmark datasets are: All included species with strain-level diversity (Supplementary Fig. D1), to explore its’ effect on program performances. They also included viruses, plasmids and other circular elements, to assess their impact on program performances. All datasets furthermore included genomes at different evolutionary distances to those in reference databases, to explore their effect on taxonomic binning. For every individual metagenome sample and for the pooled data set samples, gold standard assemblies, genome bin and taxon bin assignments, as well as taxonomic profiles were generated (available at https://data.cami-challenge.org/participate). The data generation pipeline is available on GitHub and as a docker container at https://hub.docker.com/r/cami/emsep/.

### Challenge Organization

The first CAMI challenge benchmarked software for sequence assembly, taxonomic profiling and (taxonomic) binning. To allow developers to familiarize themselves with the data types, biobox-containers and in- and output formats, we provided simulated datasets from public data together with a standard of truth before the start of the challenge (Supplementary Figs M1, M2, https://data.cami-challenge.org/). Reference datasets of RefSeq, NCBI bacterial genomes, SILVA^53^, and the NCBI taxonomy from 04/30/2014 were prepared for taxonomic binning and profiling tools, to allow performance comparisons for reference-based tools based on the same reference datasets. For future benchmarking of reference-based programs with the challenge datasets, it will be important to use these reference datasets, as the challenge data have subsequently become part of public reference data collections.

The CAMI challenge started on 03/27/2015. Challenge participants had to register on the website for download of the challenge datasets, with 40 teams registered at that time. They could then submit their predictions for all datasets or individual samples thereof. Optionally, they could provide an executable biobox implementing their software together with specifications of parameter settings and reference databases used. Submissions of assembly results were accepted until 05/20/2015. Subsequently, a gold standard assembly was provided for all datasets and samples, which was suggested as input for taxonomic binning and profiling. This includes all genomic regions from the genome reference sequences and circular elements covered by at least one read in the pooled metagenome datasets or individual samples (Supplementary Methods, Section 1.1.3). Provision of this assembly gold standard allowed us to decouple the performance analyses of binning and profiling tools from assembly performance. Developers could submit their binning and profiling results until 07/18/2015. Overall, 215 submissions representing 25 different programs were obtained for the three challenge datasets and samples, from initially 19 external teams and CAMI developers, with 16 teams consenting to publish (Supplementary Table 1). The genome data used to generate the simulated datasets was kept confidential until the end of the challenge and then released^9^. The CAMI challenge and toy datasets including the gold standard are available for download and in the CAMI benchmarking platform, where further predictions can be submitted and a range of metrics calculated for benchmarking (https://data.camichallenge.org/participate).

## ACKNOWLEDGEMENTS

We thank C. Della Beffa, J. Alneberg, D. Huson, and P. Grupp for their inputs and the Isaac Newton Institute for Mathematical Sciences for its hospitality during the programme MTG, which was supported by EPSRC Grant Number EP/K032208/1. The sequencing work conducted by the U.S. Department of Energy Joint Genome Institute, a DOE Office of Science User Facility, is supported under Contract No. DE-AC02–05CH11231. R.G.O. acknowledges support by the “Cluster of Excellence on Plant Sciences” program funded by the “Deutsche Forschungsgemeinschaft“. P.D.B was supported by the National Science Foundation under Grant No. DBI-1458689. This work used the Extreme Science and Engineering Discovery Environment (XSEDE), which is supported by National Science Foundation grant number OCI-1053575. Specifically, it used the Bridges and Blacklight systems, which are supported by NSF award numbers ACI-1445606 and ACI-1041726, respectively, at the Pittsburgh Supercomputing Center (PSC).

